# Temporal dynamics of Bacteria, Archaea and protists in equatorial coastal waters

**DOI:** 10.1101/658278

**Authors:** Caroline Chénard, Winona Wijaya, Daniel Vaulot, Adriana Lopes dos Santos, Patrick Martin, Avneet Kaur, Federico M Lauro

## Abstract

Singapore, an equatorial island in South East Asia, is influenced by a bi-annual reversal of wind directions which defines two monsoon seasons. We characterized the dynamics of the microbial communities of Singapore coastal waters by collecting monthly samples between February 2017 and July 2018 at four sites located across two straits with different trophic status, and sequencing the V6-V8 region of the small sub-unit ribosomal RNA gene (rRNA gene) of Bacteria, Archaea, and Eukaryota. Johor Strait, which is subjected to wider environmental fluctuations from anthropogenic activities, presented a higher abundance of copiotrophic microbes, including Cellvibrionales and Rhodobacterales. The mesotrophic Singapore Strait, where the seasonal variability is caused by changes in the oceanographic conditions, harboured a higher proportion of typically marine microbe groups such as Synechococcales, Nitrosupumilales, SAR11, SAR86, Marine Group II Archaea and Radiolaria. In addition, we observed seasonal variability of the microbial communities in the Singapore Strait, which was possibly influenced by the alternating monsoon regime, while no seasonal pattern was detected in the Johor Strait.

## Introduction

Marine planktonic communities harbour representatives from all three domains of life (Bacteria, Archaea and Eukaryota). Together, these organisms perform a range of global biological and geochemical processes. In contrast to the open ocean, coastal environments are influenced by local disturbances such as freshwater land run-off, land to sea transfer of nutrients and organic matter through river discharge and mixing in shallow areas induced by tidal currents, in addition to seasonal cycles. These variable conditions may lead to complex dynamics where patterns of reoccurring microbial communities are harder to access and predict.

In temperate waters, annual cycles of plankton communities have been studied for a long time^1, 2^. The seasonal changes in microbial community composition and biomass are often attributed to responses to changes in environmental conditions driven by seasonal climate cycles^3–6^. For example, Lambert *et al.*^4^ demonstrated a strong rhythmicity in Bacteria, Archaea and protist communities despite sporadic meteorological events and irregular nutrient availability in a coastal site situated in the North Western Mediterranean Sea. In contrast, few studies have investigated the seasonal dynamics of coastal microbial communities in equatorial waters.

Equatorial waters are subjected to monsoons, periods when the prevailing winds over land and adjacent ocean areas reverse directions on a seasonal basis in response to differences of heating pattern between land and ocean. These differences alter the patterns of rising and sinking air near the equator, resulting in the seasonal migration of the intertropical convergence zone (ITCZ), an area of low atmospheric pressure where Northeast and Southeast Trade Winds meet^7^. By influencing vertical mixing^8^, upwelling^9^ and advective transport^10^, monsoon systems have previously been shown to influence phytoplankton dynamics in tropical waters^8, 11, 12^. Miki *et al.*^8^ demonstrated a higher concentration of chlorophyll during the NE monsoon than the SW monsoon in the Sulu Sea off the southwest side of the Philippines. A similar trend in chlorophyll concentration was also observed in the Strait of Malacca^11^.

In Singapore, a highly urbanized island city state of South East Asia, located just one degree North of the equator, a bi-annual reversal of wind directions and two monsoon seasons named after the prevailing wind direction^7^ characterize the annual meteorological conditions in the island. The Northeast Monsoon (NE Monsoon), generally occurring from December to early March, brings primarily north-easterly winds and long periods of heavy rain across the region, while the Southwest Monsoon (SW Monsoon), from June to September, exhibits south-westerly winds with higher temperatures, scattered showers and thunderstorms mostly occurring in the afternoon. Inter-monsoon periods are generally less windy and experience lower precipitation. Singapore is also surrounded by two main bodies of water with different trophic status: the Singapore Strait in the South and the Johor Strait in the North. The Singapore Strait exhibits oceanic conditions and is influenced by strong tidal currents up to 2 m s^−1^ and alternate oceanic influxes from the South China Sea and the Java Sea^13^. The Johor Strait, less than 1 km wide, is subjected to greater environmental fluctuations resulting from a combination of anthropogenic activities, sporadic riverine inputs and reduced tidal mixing^14^. Recurrent spring/summer blooms have been reported during the inter-monsoon months April/May in the years of 1935, 1948, 1949 and 1968^15^. More recent studies either applied morphological identification methods to a restricted number of taxa^14^ or used techniques such as flow cytometry and pigments analysis that have limited taxonomic resolution^16, 17^ to investigate the dynamics of the plankton communities in Singapore coastal waters.

Here, we present an 18-month study that uses SSU rRNA metabarcoding to characterize the temporal variation of the marine microbial community in Singapore waters. We observed taxonomic compositions reflecting differences in trophic status between the two Straits and identified a seasonal signature in the microbial community composition of the Singapore Strait, probably linked to the monsoonal current reversal. In contrast, no seasonal patterns, but large month-to-month changes, in beta-diversity were observed in the Johor Strait, suggesting that localized pulse disturbances might be responsible for shaping community composition.

## Materials and Methods

### Sampling sites and water collection

Surface water (~ 1 m depth) was collected with a submersible pump monthly from February 2017 to July 2018 at four different stations around Singapore (Figure S1, Supplementary Data S1). Sampling was performed within two days of neap tide, except for samples # 21 collected during the full moon (28 June 2018). Two stations were in the Singapore Strait, St. John (1.2383° N, 103.8536° E) and East Coast (1.2972° N, 103.9207° E), and two in the Johor Strait, Pasir Ris (1.3886° N, 103.9515° E) and Sembawang (1.4716° N, 103.8051° E). Four monsoon periods were identified using the prevailing wind direction from the annual climatological report for 2017 and 2018 from the Meteorological Service of Singapore^18^: Northeast (NE), inter-monsoon 1 (IM-1), Southwest (SW), and inter-monsoon 2 (IM-2). A month was assigned to the NE or SW Monsoon if the wind recorded at Changi airport blew for more than 50% of the month from either the first quadrant (NE) or third quadrant (SW) direction, respectively, and assigned to the IM periods in all other cases.

Temperature and salinity were measured at ~ 1m using a Eureka Water Probes Manta+ Trimeter Multiprobe. Chlorophyll was measured using an AquaFlash™ Handheld Active Fluo-rometer (Turner Designs). For dissolved inorganic nutrients, water was syringe-filtered (0.22 *µ*m Acrodisc, PALL) into acid-washed, polypropylene centrifuge tubes, immediately flash-frozen in liquid nitrogen, and stored at −20° C until analysis. For microbial community samples, 1L of seawater was first filtered through a 150 *µ*m net mesh filter to remove large detritus and zooplankton, and then the biomass was collected on 0.22 *µ*m Sterivex filters (Millipore). The Sterivex filters were then immediately stored at −80° C until further analysis. Samples for flow cytometry (980 *µ*L) were fixed for 15 min at 4° C in the dark with 25% electron microscopy (EM) grade glutaraldehyde (0.5% final concentration; Sigma-Aldrich), flash-frozen in liquid nitrogen and stored at −80° C until analysis.

### Nutrient analysis

Samples for dissolved inorganic macronutrients, NO_3_+NO_2_ (referred to as NO_x_ hereafter), NO_2_, NH_4_, PO_4_, and Si(OH)_4_ were thawed at room temperature and immediately measured on a SEAL AA3 segmented-flow autoanalyser according to SEAL methods for seawater analysis. NH_4_ was measured fluorometrically^19^. For samples collected on 3 April 2017, 02 May 2017 and 31 May 2017, the dissolved inorganic macronutrients were measured using the APHA^20^ method and Si (4500-SiO_2_ Flow injection analysis for Molybdate-Reactive Silicate) as a service provided by DHI-Singapore Seawater. Total dissolved inorganic nitrogen (DIN) was calculated as NO_x_+NH_4_.

### Enumeration of bacteria

Bacteria were enumerated in duplicate water samples by flow cytometry using a CytoFLEX (Beckman Coulter, Singapore) equipped with blue (488 nm) and violet (405 nm) lasers. Prior to analysis, samples were thawed and diluted (between 5-10x) with 0.2 *µ*m filtered sterile 10 mM Tris, 1 mM ethylene-diamine-tetra-acetic acid buffer (pH 8.0) and stained with SYBR Gold (Life Technologies, Singapore). Samples were incubated in the dark for 15 minutes prior to analysis. The analysis was performed for 1 minute at medium flow rate (~ 30 *µ*L min^−1^) with an event rate of ~ 200-1200 particles per second with a threshold set on the green fluorescence channel (525/40 BP). Bacteria were discriminated based on their signature on a scatter plot of green fluorescence against violet side-scatter using CytExpert software provided with the CytoFLEX.

### DNA extraction and 16S/18S PCR amplification and sequencing

DNA was extracted from the Sterivex filters using a modified protocol for MoBio PowerSoil kit (MoBio Laboratories, Carlsbad, CA), as previously published by Jacobs *et al.* 2009^21^. Briefly, the Sterivex filter casing was broken open and the filter was cut with a sterile razor blade into small lengthwise strips and placed into the PowerBead tube. 60 *µ*L of the manufacturer solution C1 were added to the tube which was incubated for 10 min at 70° C with constant shaking (500 rpm) twice and vortexed at maximum speed between each incubation. Following the last incubation, 700 *µ*L of phenol-chloroform-isoamyl alcohol was added into the PowerBead tube and vortexed at maximum speed for 10 min. The PowerBead tube was centrifuged at 10,000 g for 30 seconds and 800 *µ*L of supernatant transferred to a new collection tube to perform the DNA extraction as per the manufacturer’s instructions. DNA was eluted into 100*µ*L of nuclease-free water and quantified using a Qubit® 2.0 fluorometer with the dsDNA Broad Range assay kit (Invitrogen, Singapore). Amplicon libraries were then generated using a modified version of the Illumina 16S Metagenomic Sequencing Library Preparation Protocol^22^. Universal primer pairs (926wF: 5’-AAACTYAAAKGAATTGRCGG-3’ and 1392R: 5’-ACGGGCGGTGTGTRC-3’) targeting the V6-V8 hyper-variable region of the 16S/18S Ribosomal RNA gene with an Illumina-specific overhang were used^23^. For each sample, triplicate PCR reactions obtained with 22 cycles were pooled and purified using Agencourt AMpure XP beads (Beckman Coulter, Singapore). Products from this first step were then sent to the sequencing facility at the Singapore Centre for Life Science and Engineering (SCELSE) where a second round of PCR was performed to add dual barcodes to each amplicon library. Afterwards, PCR products were purified with Agencourt AMpure XP beads, pooled at equal volume and sequenced with an Illumina MiSeq sequencing run (2×300 bp).

### Sequence analysis

The raw sequencing data was initially processed by removing primers with cutadapt^24^. Paired-end joining, denoising and taxonomic assignment of Amplicon Sequence Variants (ASV) were performed using Quantitative Insights Into Microbial Ecology (QIIME) release 2018.6^25^. Briefly, after importing the sequence data into the QIIME2 environment, the denoising and pair-end joining were performed using DADA2^26^. After discarding all the ASVs with less than 10 sequence representatives, phylogenetic relationships were inferred by aligning representative sequences with MAFFT^27^, filtering the alignment and constructing phylogenetic trees using fasttree^28^ with a midpoint root. Taxonomic classification was performed using the classify-sklearn method^29^ using the SILVA version 132 database as a reference. All ASVs identified as eukaryotic (nuclear, plastid) were further assigned against the PR^2^ database^30^ version 4.12.0 (https://github.com/pr2database/pr2database) using the RDP naive Bayesian classifier^9^ as implemented in the R *dada2* package. ASVs with a bootstrap value lower than 90% at the superegroup level were discarded. For statistical analyses, Metazoa (mostly copepods) were removed from the final ASV table because they are likely to represent eggs or organismal fragments that were not captured by the pre-filtering step. Similarly, eukaryotic plastid sequences were removed to avoid counting photosynthetic organisms twice. The final ASV table contained a total of 2,571 ASVs.

### Statistical analyses

Pairwise comparisons between the different locations (Johor Strait, Singapore Strait) and monsoon periods were performed using ANOSIM with the software R^31^. Pairwise community dissimilarity was calculated using Bray-Curtis distance as implemented in the *vegan* package. The Shannon index (H’) and richness calculated with the *vegan* package, nMDS plots were generated with the metaMDS function also in the *vegan* package. The Envfit analysis generated using the *vegan* package used the following environmental parameters: salinity, temperature, chlorophyll, bacterial abundance, PO_4_, DIN, Si(OH)_4_,NH_4_ and amount of rainfall seven days (rain-7) prior to the sampling day. The differential abundance of relevant ASVs was extrapolated using Gneiss^32^, with Ward’s correlation clustering as a function of monsoon period for the Singapore Strait. The ASVs contributing significantly to the difference between the two monsoon regimes were computed using the DESeq2 package^33^ using a threshold for the corrected p-value of 0.01. The magnitude of changes occurring in the community during the time series was inferred with the first-distances method^34^ applied to the unweighted Unifrac distance matrix^35^.

## Results

### SPhysico-chemical parameters in Singapore coastal waters

Despite their proximity and physical connection, the Singapore and Johor Straits exhibit different environmental conditions (Figure 1, Figure S2, Supplementary Data S1). Temperature showed little variation, but was slightly higher in the Johor (29.0–32.8°C) than the Singapore (27.1–30.6°C) Strait (Figure S2A). The lowest temperatures were consistently found during the NE monsoon. Much larger variability was observed in salinity: the two sites in the Singapore Strait were very similar to each other, with salinity ranging from 29.8 to 32.6 and lowest values during the SW monsoon (Figure 1). In contrast, salinity was lower and more variable in the Johor Strait, ranging from 18.1 to 30.3, with lowest values mostly during the NE monsoon. Sembawang, the sampling site closest to the causeway between Singapore and Malaysia, typically had lower salinity than Pasir Ris (Figure 1).

**Figure 1.**
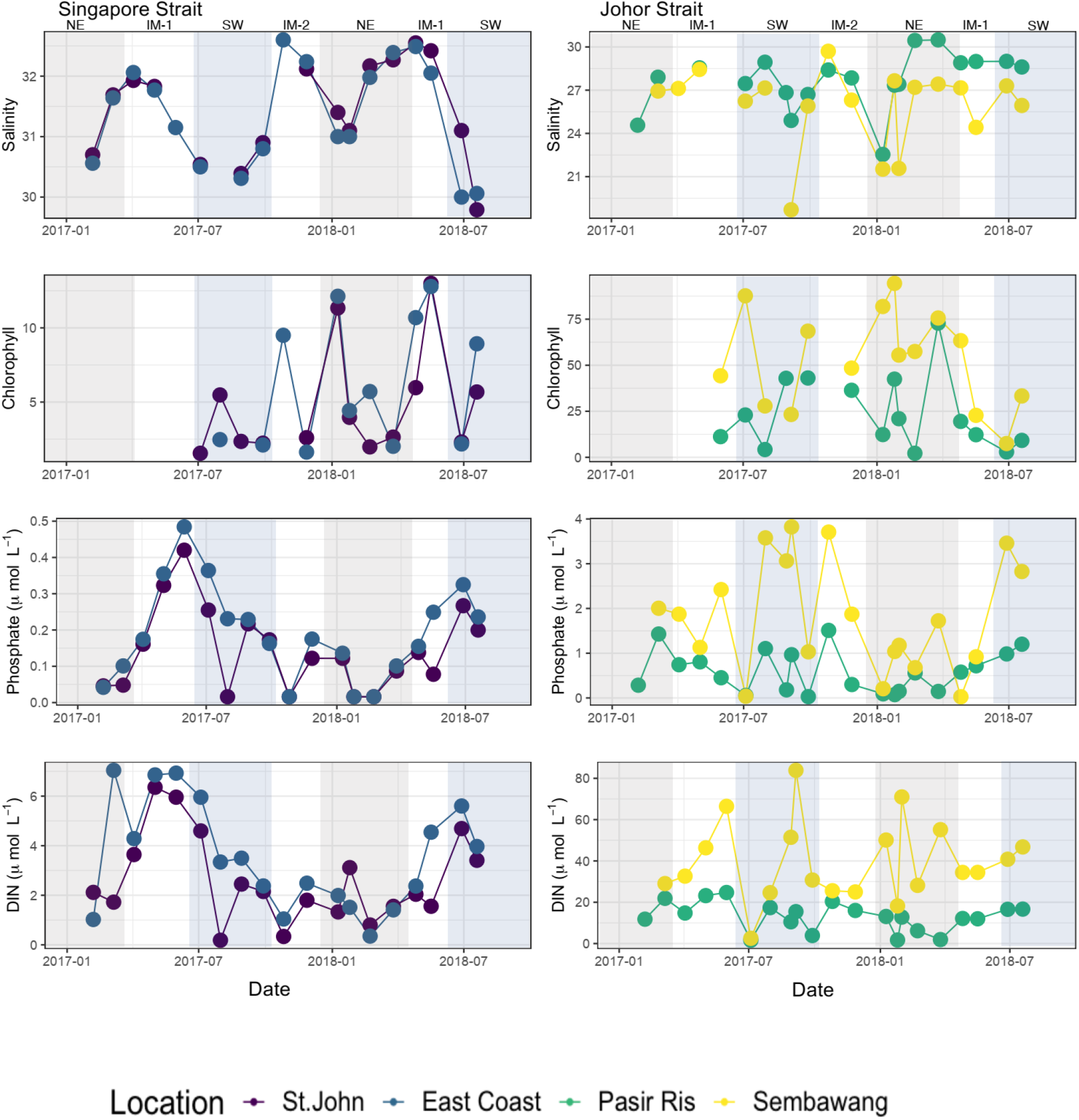
Salinity, chlorophyll *a*, phosphates and DIN (dissolved inorganic nitrogen) during the 18-month time series in Singapore coastal waters. Highlights in grey and blue represent NE and SW monsoon, respectively.

Dissolved nutrient concentrations also differed strongly between the Singapore and Johor Straits: NO_x_, NO_2_, NH_4_ and PO_4_ were typically 2–10-fold higher in the Johor compared to the Singapore Strait, while Si(OH)_4_ was on average more similar, but with higher variability in the Johor Strait (Figure 1, Supplementary Data S1, Figure S2). The Singapore Strait showed more oligo-to mesotrophic characteristics, with NO_x_ ranging from < 1–5 *µ*mol L^−1^, NO_2_ and NH_4_ almost always < 0.5 *µ*mol L^−1^ and often close to or below detection limits, and PO_4_ frequently < 0.2 *µ*mol L^−1^. Si(OH)_4_ in the Singapore Strait showed a seasonal pattern inverse to that of salinity. In contrast, the Johor Strait mostly had NO_x_ > 5 *µ*mol L^−1^ and often > 10 *µ*mol L^−1^, with NO_2_ regularly contributing more than 50% of NO_x_. NH_4_ mostly ranged at Pasir Ris from 1 to 8 *µ*mol L^−1^, and at Sembawang mostly around 10–40 *µ*mol L^−1^, reaching as high as 75 *µ*mol L^−1^. PO_4_ in the Johor Strait was generally in the range of 0.5–3.0 *µ*mol L^−1^, but occasionally dropped to < 0.1 *µ*mol L^−1^. Higher values were usually found during the SW monsoon. In the Singapore Strait, NO_x_ and PO_4_ were generally highest just before or during the SW monsoon, and lowest during the NE monsoon. The Johor Strait displayed less coherent seasonal patterns, with NO_x_ and PO_4_ showing opposite seasonal trends in Sembawang, and no clear pattern for Si(OH)_4_. There were no differences in total precipitation for the 7 days prior to sampling between the Johor and Singapore Straits throughout the year, with values ranging from 0 to 128 mm (Supplementary Data S1).

Bacterial abundance was different between the Singapore and Johor Straits (Supplementary Data S1, Figure S2), with abundances ranging from 0.4×10^6^ to 1.6×10^6^ cells*⋅*mL^−1^ and 1.1×10^6^ to 9.4×10^6^ cells*⋅*mL^−1^, respectively. In general, higher bacterial counts were recorded at Sembawang.

Chlorophyll concentration varied substantially between sites, broadly reflecting the concentration of nutrients: low values were found in the Singapore Strait (mostly <2 *µ*g L^−1^, but up to 13 *µ*g L^−1^), while the Johor Strait rarely had less than 10 *µ*g L^−1^, and up to 94 *µ*g L^−1^ (Figure 1, Supplementary Data S1). In the Singapore Strait, chlorophyll was lowest during the SW monsoon and highest during inter-monsoon 1 or the NE monsoon, while in the Johor Strait, concentrations were highest during the NE monsoon or inter-monsoon 2, and lower during the SW monsoon.

### Community composition of Singapore coastal waters

A total of 70 samples, 30 from the Singapore Strait and 40 from the Johor Strait from February 2017 to July 2018 were analyzed by rRNA metabarcoding using primers that amplify both prokaryotic 16S and eukaryotic 18S rRNA^23^. After quality control filtering, end-pair joining and chimera filtering, a total of 5,036,783 sequences were retained with an average of 71,954 sequences per sample which were assigned to 2,556 ASVs (Supplementary Data S2).

Based on the Shannon index (H’), Singapore Strait communities generally had a higher diversity than those from the Johor Strait for Archaea and Eukaryota, but not for the Bacteria (Figure 2). The richness, inferred from the number of ASVs, was also generally higher in the Singapore Strait for Archaea and Eukaryota (Figure 2).

**Figure 2.**
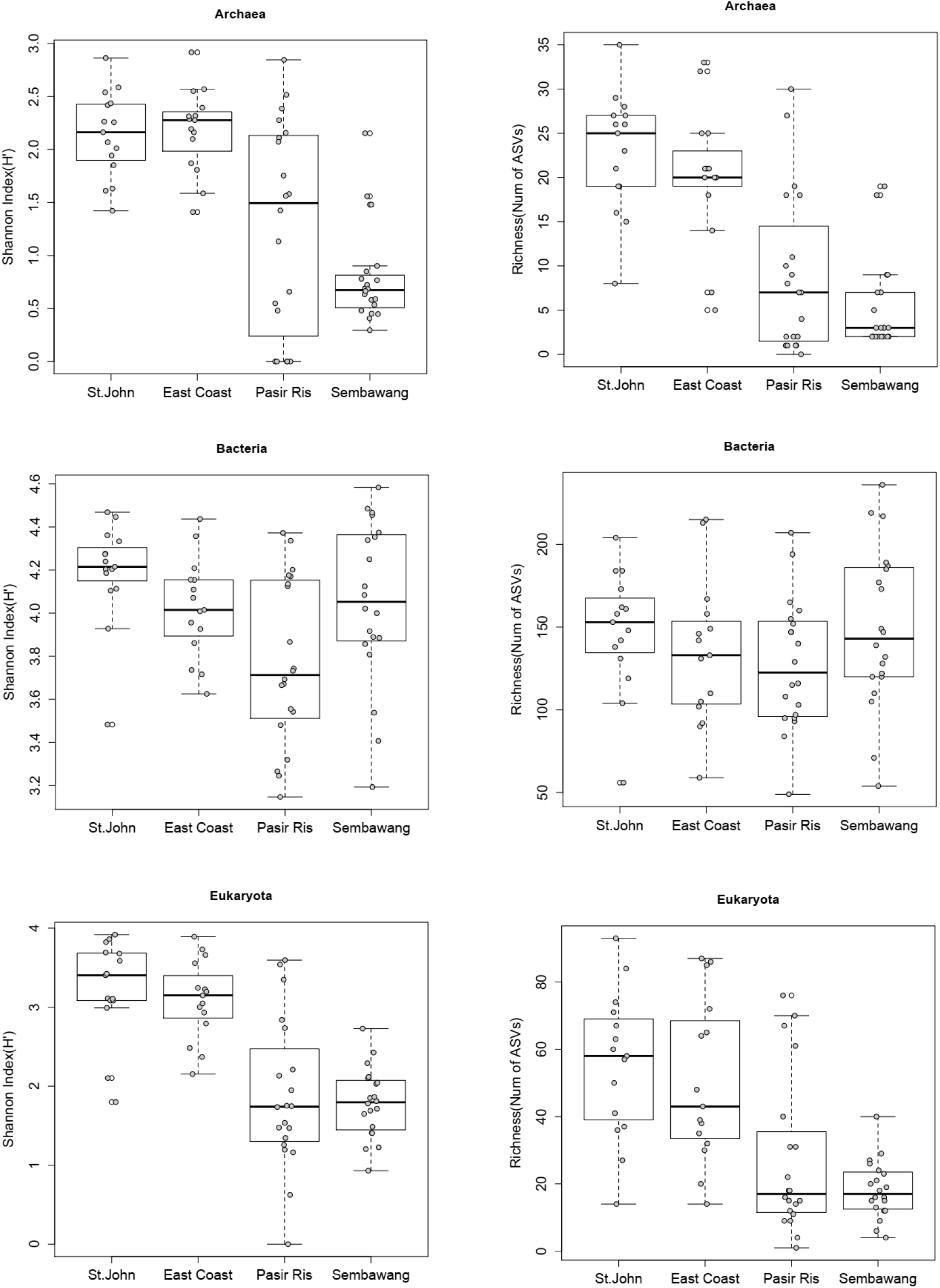
Shannon Index (H’, left) and Richness (Number of ASVs, right) of microbial communities at 4 stations in Singapore coastal waters for Archaea (top,A-B), Bacteria (middle,C-D) and Eukaryota (bottom, E-F). Empty dots represent outliers.

The taxonomic composition of Bacteria, Archaea and Eukaryota communities varied between straits (Figures 3, S3 and S4). At the domain level, Bacteria dominated both straits but Archaea were more prevalent than Eukaryota in the Singapore Strait than the Johor Strait (Figure 3).

**Figure 3.**
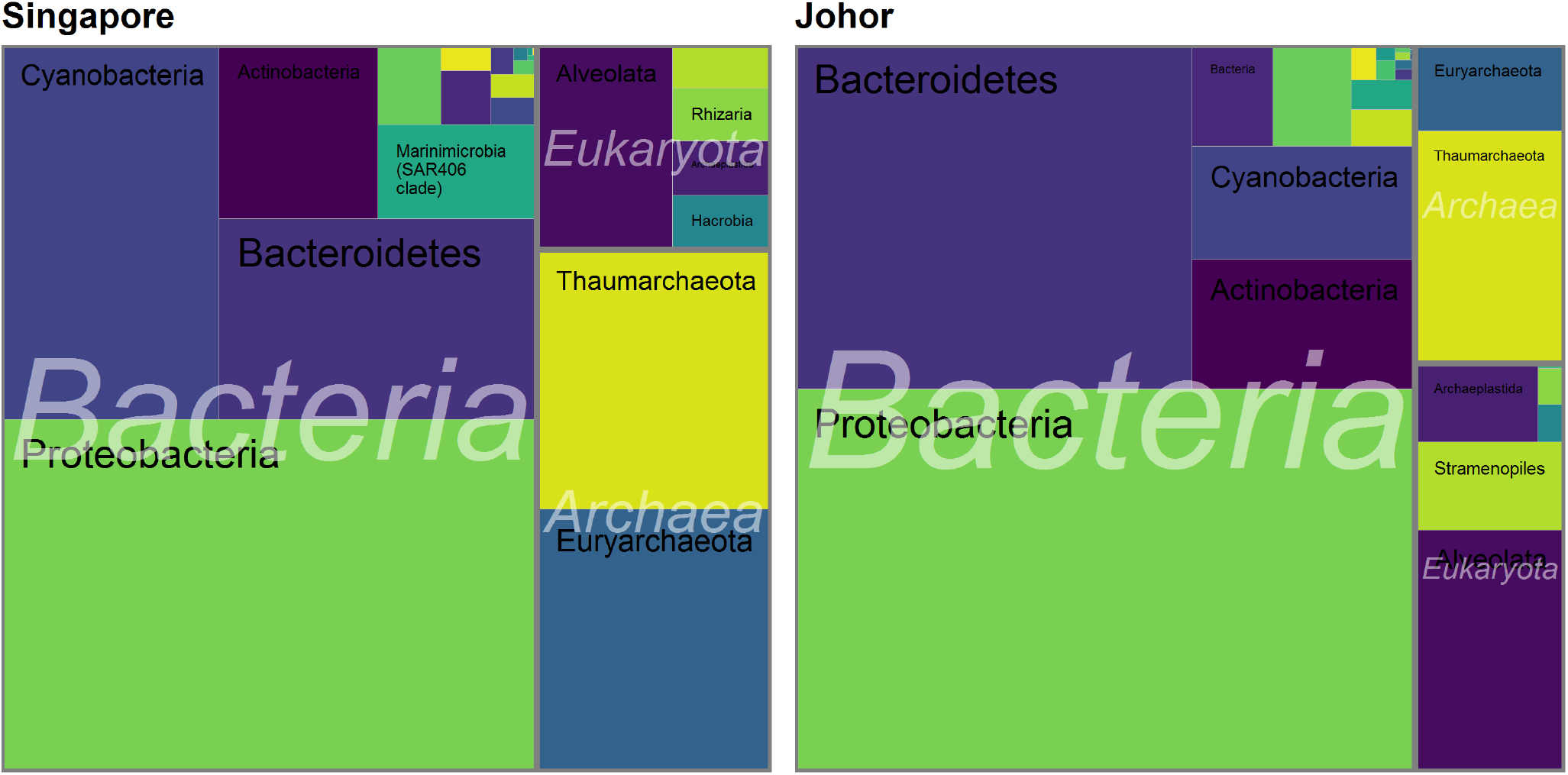
Tree map with the major taxonomic groups in Singapore and Johor Straits.

Among prokaryotes, Cyanobacteria, specific proteobacterial groups and the archaeal phyla Euryarchaeota and Thaumarchaeota were over-represented in the Singapore Strait. Within the Archaea, the phylum Thaumarchaeota was mainly represented by the order Nitrosupumilales, which was present in both the Singapore and Johor Straits, but had a higher relative proportion in the Singapore Strait (Figure S4). In the Singapore Strait, both *Candidatus* Nitrosopelagicus and *Candidatus* Nitrosopumilus were abundant while *Candidatus* Nitrosopumilus was abundant in the Johor Strait (Figure S5. Euryarchaeota were mostly found in the Singapore Strait, with many ASVs assigned to marine group II Euryarchaeota (MGII). Marine group III Euryarchaeota (MG-III) were also present in the Singapore Strait, but in lower proportion, and absent in most samples from the Johor Strait. Other prevalent members of the microbial communities of the Singapore Strait included the alphaproteobacterial group SAR11, the gammaproteobacterial group SAR86, the deltaproteobacterial group SAR324 and the candidate actinobacterial genus *Actinomarina*. In the Johor Strait, Proteobacteria and Bacteroidetes were the most abundant groups (Figure S4). The most abundant family within the Bacteroidetes in Johor included the Flavobactericeae and the NS5 marine group. Within the Proteobacteria there was a high abundance of reads assigned to *Roseobacter* strain HIMB11, to members of the gammaproteobacterial OM60/NOR5 clade, Cellvibrionales (in particular members of the genus *Luminiphilus*) and the family Burkholderiaceae, Figure S4, Figure S3, Figure S6).

The eukaryotic planktonic community in both straits was dominated by Dinoflagellata (Alveolata), Ochrophyta (Stramenopiles, mostly diatoms) and Chlorophyta (Figure S4-C). Among Dinoflagellata, Dinophyceae and Syndiniales were the main groups in the Singapore Strait, while the Johor Strait was dominated by Dinophyceae (Figure S4-C). Most of Dinophyceae reads from the Johor Strait were affiliated with the order Gymnodiniales (e.g. *Gyrodinium* and *Woloszynskia*) and Gonyaulacales, while the Singapore Strait reads were affiliated with uncultured dinoflagellate sequences (Figure S5). For example, the sequence of the most abundant Dinophyceae ASV (0115) in the Singapore Strait is 100% similar to that of a marine eukaryote clone o10.5-16 (KX532243) from the South China Sea (Supplementary Data S2). The Dinophyceae community in the Singapore Strait was more diverse than in the Johor Strait, where two ASVs (0011, *Gyrodinium* and 0043, *Gonyaulax*) practically dominated the whole community (Figure S5). These ASVs were also within the top 50 most abundant ASV when considering the whole dataset (Bacteria, Eukaryota and Archaea).

Reads assigned to the marine parasites Syndiniales were nearly absent in the Johor Strait. In fact, the most abundant Syndiniales ASV found in the Singapore strait were absent in Johor (Figure S5 Supplementary Data S2). Syndiniales are divided into 5 main groups^36^. Syndiniales sequences from the Singapore Strait were mainly affiliated to the highly diverse group II (Figure S7), which contains a single genus formally described, *Amoebophrya*, which infects a wide range of dinoflagellate hosts. Most Syndiniales reads could only be assigned to uncultured clades known from environmental sequences (Supplementary Data S2). For example, the sequence of ASV-0285, the main Syndiniales ASV from the Singapore Strait, was 100% similar to the uncultured eukaryote clone ST5900.074 (KF130025) obtained from the South China Sea.

Among the other Alveolata groups, very few reads were assigned to Apicomplexa (e.g Gregarines and Perskinsea, Supplementary Data S2). Only Ciliophora were found in both straits, but more abundant in the Singapore Strait (Figure S4-C). Ciliophora reads were mainly assigned to the class Spirotrichea, with the genera *Parastrombidinopsis* and *Pelagostrobilidium* present in the Johor Strait *vs. Strombidiida* and the *incertae sedis* genus *Mesodinium* in the Singapore Strait (Figure S5).

Bacillariophyta (Ochrophyta) was the second and third most abundant group in the Johor and the Singapore Straits, respectively (Figure S4). Mediophyceae (polar centrics) were well represented in the two straits while Coscinodiscophyceae (radial centrics) were mainly found in the Singapore Strait. *Cyclotella*, *Cerataulina* and *Thalassiosira* (all Mediophyceae) were the main genera found in the Johor Strait, while *Leptocylindrus* (Coscinodiscophyceae), *Guinardia* (Coscinodiscophyceae) and *Skeletonema* (Mediophyceae) dominated the Singapore Strait (Figure S5). The highly diverse genus *Chaetoceros* (Mediophyceae) was the only dominant genus common to both straits. Only two ASVs with a low number of sequences were assigned to Bacillariophyceae, which contains pennate diatom genera such as the toxic blooming *Pseudonitzschia* (Supplementary Data S2). Other classes of Ochrophyta (except for ASV-2888 belonging to Raphidophyceae) were not detected based on nuclear 18S rRNA. However Pelagophycae, Dictyochophyceae and Chrysophyceae, all Orchophyta classes, were detected based on plastid 16S rRNA (Supplementary Data S2). Stramenopile groups such as the diverse group of heterotrophic flagellates MAST (Marine stramenopiles) and fungus-like members of Oomycota and Labyrinthulea were also found in both straits, although in low abundance (Figure S4-C and Supplementary Data S2).

Mamiellophyceae and Trebouxiophyceae were the two main classes of Chlorophyta (green algae) found in the Singapore and Johor Straits, respectively. Trebouxiophyceae ASVs were assigned to the highly diversified marine and brackish water coccoid genus *Picochlorum* (Figure S5), known for its broad halotolerance^37^,^38^. Members of the two widespread Mamiellophyceae genera, *Micromonas* and *Ostreococcus*, were found in both straits (Figure S7 and Supplementary Data S2). *Micromonas* was the main photosynthetic genus in the Singapore Strait (Figure S5) two clades, one corresponding to the ubiquitous species *M. commoda*, and the other, to the undescribed clade B5,^39^ found at the two stations in the Singapore Strait all year around (Figure S7). Two clades of *Ostreococcus* were found mainly at Pasir Ris station in the Johor Strait during the SW monsoon (Figure S7): Clade B, which seems to be the dominant clade in tropical waters^39^, and another clade that could be the new Clade E, which was found in Singapore during the Ocean Sampling Day survey^39^, and for which no culture is available yet. The ubiquitous genus *Bathycoccus* was less abundant in Singapore waters, appearing sporadically (Supplementary Data S2).

Within the supergroup Hacrobia, reads belonging to Haptophyta and Cryptophyta divisions were the most abundant. Among haptophytes, all ASVs belonged to the class Prymnesiophyceae (Figure S5 and Supplementary Data S2) and were closely related to the widespread and non-calcifying genera *Chrysochromulina* and *Phaeocystis*, mostly in Singapore Strait. Only ASV-1427, with low abundance, was assigned to the calcifying coccolithophorids order Isochrysidales (*Gephyrocapsa*) (Supplementary Data S2). Cryptophytes were mainly detected in the Singapore Strait and assigned to the genus *Geminigera* (Figure S5).

Reads belonging to the supergroup Rhizaria were found in both straits. Radiolaria were dominant in the Singapore Strait but nearly absent in the Johor Strait. The most abundant Radiolaria ASV was assigned to the order Taxopodia clade RAD-B (Figure S5). Rhizaria reads from the Johor Strait belonged to the division Cercozoa and were assigned to the Class Filosa-Imbricatea (Figure S5)^40^.

### Factors structuring microbial communities off Singapore

Non-metric multidimensional scaling analysis (nMDS) of microbial community composition revealed that samples clustered based on strait (Figure 4). While there was little dissimilarity between the St. John and East Coast sites of the Singapore Strait cluster, there was a clear separation between the Johor Strait sampling sites (Sembawang and Pasir Ris stations). An analysis of similarity (ANOSIM) based on different grouping factors (strait, site, monsoon) confirmed significant differences based on strait and site (Table 1). Envfit analysis revealed that most environmental factors (salinity, temperature, chlorophyll *a*, DIN, NH_4_ and bacterial abundance were significant drivers (p < 0.05) for the difference between straits (Figure 4).

**Table 1.** ANOSIM analysis.

| Sample set | Grouping | R | Significance |
| --- | --- | --- | --- |
| All sites | Strait | 0.802 | 0.001 |
|  | Site | 0.652 | 0.001 |
|  | Monsoon | 0.005 | 0.34 |
| Singapore Strait | Site | -0.072 | 0.498 |
|  | Monsoon | 0.270 | 0.002 |
| Johor Strait | Site | 0.421 | 0.001 |
|  | Monsoon | -0.026 | 0.675 |

**Figure 4.**
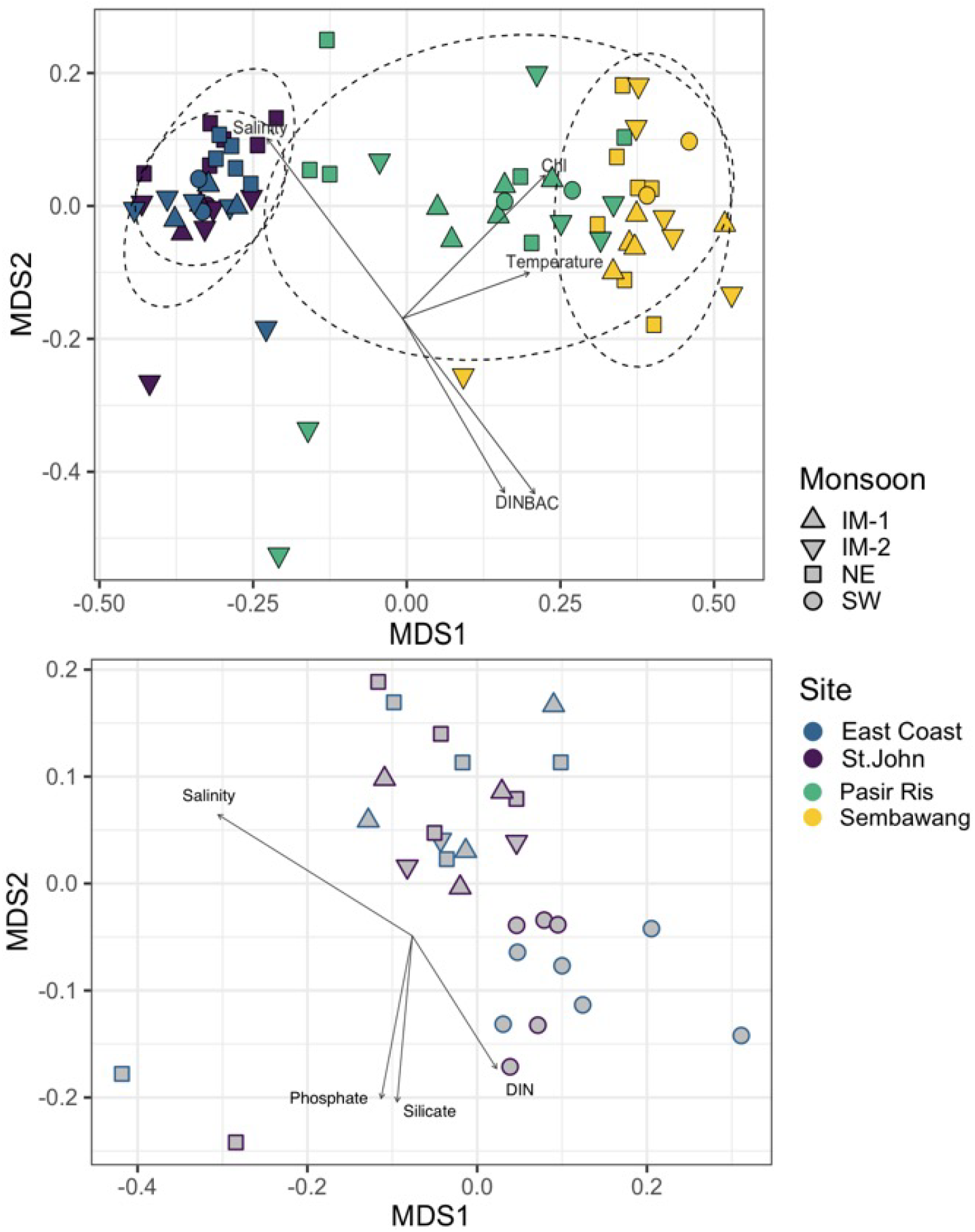
Non-metric multidimensional scaling (nMDS) analysis of Bray-Curtis similarity index. Each sample is labelled by location and monsoon period. Arrows represent environmental parameters with p < 0.001 when performing an *envfit* analysis The ellipses represent 95% confidence intervals for samples collected at the same station. Top. All stations. Bottom. Only Singapore Strait samples (STJ and EC stations).

An nMDS analysis taking only into account the Singapore Strait displayed little overlap between the SW monsoon and the others monsoon periods (Figure 4). ANOSIM confirmed that there was a significant difference based on monsoon (Table 1) and this was confirmed by gneiss balance analysis (Figure S9-top). PO_4_, DIN and Si(OH)_4_ were the most significant environmental variables influencing separation of SW-Monsoon cluster from other monsoon periods (p < 0.1, Figure 4). In contrast, the nMDS restricted to Johor Strait samples did not show clustering based on monsoon (Figure S8) and ANOSIM demonstrated a significant difference only based on site (Table 1).

### Seasonal variability in Singapore coastal water communities

Seasonal variability associated with the monsoon was observed in the Singapore Strait. Linear regression models implemented in Gneiss analysis indicated a significant difference between SW and NE monsoon regime (Figure S9top), which was driven by an increase in abundance of members of the order Nitrosopumilaceae (Figure S6.

A DESeq2 analysis revealed that the abundance of 71 specific taxa was influenced by the NE or SW monsoon. Among those ASVs related to Alteromonadales (0292, 0512, 0747), Pirellulales(0514, 0490), SAR11(0217, 0407, 0448, 0130, 0181, 0280, 0074, 0054) SAR116(0371) and *Synechococcus* clade CRD1 (0277) were associated with the SW monsoon, while the ASVs related to Desulforculaceae (0772, 0921), Opitutales MB11C04 marine group (0819, 0274) and *Cyanobium* (0424, 0470) were associated with the NE monsoon. For the archaea, different ASVs assigned to the Marine Group II and Marine Group III showed a change in their abundance by monsoon based on the DESeq2 analysis. ASVs related to Marine Group II (0398, 0185, 0358, 0101) and Marine Group III (0297) were more abundant during the SW monsoon while ASVs related to Marine Group II (0403,441) and Marine Group III (0423) were more abundant during the NE monsoon. An abundant archaeal ASV (0005) assigned to *Candidatus* Nitrosopumilus also showed an increase during the SW monsoon. Among eukaryotes, thirteen ASVs had their abundance influenced by the monsoon according to DESeq2 analysis (Figure S9). ASVs assigned to Syndiniales group I (clades 1 and 2) and group II (clades 7 and 10-11), Mediophyceae and Ciliophora genus *Mesodinium* had their abundance increased or detected during the SW monsoon (Figure S9). During the NE monsoon an increase in abundance of ASVs related to the Syndiniales group I (clades 1 and 4) and II (clade 16), Dynophyceae order Gymnodiniales, Coscinodiscophyceae genus *Guinardia* and Ciliophora Class Spirotrichea was observed. Within these 13 ASVs, only 3 (0532, 0380 and 0324) were among the top 30 most abundant ASVs in the Singapore Strait (Figure S5).

Volatility values for the first distance analysis of the unweighted Unifrac matrix (Figure S10), which can be interpreted as the month-to-month change in the composition of microbial communities, were generally higher for samples from the Johor Strait (Sembawang and Pasir Ris) and lower for the Singapore Strait sites (St. John and East Coast). The comparison of Figure 1 and Figure S10 revealed that, in the Johor Strait, peaks in volatility corresponded to large month-to-month variations in salinity.

## Discussion

### Johor Strait microbial community is influenced by short term events

The lower values and the absence of a clearly defined seasonal pattern in salinity compared to the Singapore Strait (Figure 1) suggest that Johor Strait is influenced by river and land runoff. The lower salinity at Sembawang compared to Pasir Ris also suggests that the central portion of the Johor Strait is affected by land-runoff from storm drains, reservoirs and smaller rivers such as the Sungai Tebrau, located just north of Sembawang station. The oscillation in salinity was also reflected in the eukaryotic community composition, which harbored brackish water (e.g *Cyclotella*) and marine euryhaline groups (e.g *Picochlorum*). Groups known to have an exclusive marine lifestyle such as Syndiniales and *Micromonas* were absent at Sembawang samples and only sporadically found in the samples from Pasir Ris (Figure S7). On the other hand, the cosmopolitan genus *Gyrodinium*, known to inhabit marine, brackish and freshwaters, was found at both Johor stations (Figure S7).

The Johor Strait is clearly eutrophic compared to the Singapore Strait, with consistently higher nutrient and chlorophyll concentrations. Higher concentrations of NO_2_ and NH_3_, and the higher bacterial counts, indicate that heterotrophic recycling is likely more pronounced in the Johor Strait, consistent with eutrophication. It is unclear to what extent the nutrient concentrations in the Johor Strait are controlled by direct inputs *via* run-off, or whether sedimentary recycling processes are significant. The fact that the concentration of DIN is inversely correlated to salinity in the Johor Strait (Figure S11) but not to precipitation (data not shown) suggests that reservoir run-off processes may play an important role. Moreover, the Johor Strait exhibited a higher abundance of copiotrophic families of microbes such as the Flavobactericeae, the Burkholderiaceae, taxa affiliated with the *Roseobacter* and the OM60/NOR5 clades. All these groups have been previously implicated in nutrient remineralization and rapid responses to pulses of organic carbon such as those resulting from phytoplankton blooms and coastal runoff^41, 42^. The most abundant prokaryotic ASV in the Johor Strait *Roseobacter* strain HIMB11 (Figure S5 was also previously shown to have the genomic potential for degradation of algal-derived compound such as DMSP^43^. The Burkholderiaceae have also been shown to be one of the dominant groups in the network of urban waterways of Singapore^44^ which might represent a reservoir and source for their presence in Johor Strait.

A lower alpha diversity and a larger beta diversity was observed in the Johor compared to the Singapore Strait. The overall lower alpha diversity in the Johor Strait was mostly driven by Eukaryota and Archaea. The frequent and short pulse disturbances (short-term events with release of nutrients) experienced by the Johor Strait planktonic community seemed to affect its stability and ultimately the diversity of the system^45^. The wider Johor sample dispersion in the nMDS plots suggested little mixing between the planktonic communities from the Johor and the Singapore straits as well as within the Johor Strait, as supported by the cluster separation between Pasir Ris and Sembawang samples (Figure 4). The seasonal change in regional seawater circulation due to monsoon wind reversal did not seem to affect the microbial community composition of the Johor Strait directly. The only seasonal trend we observed in the community, linked to the SW monsoon, was the presence of the two pico-phytoplanktonic genera *Ostreococcus* and *Micromonas* (clade B5) in Pasir Ris samples during July of 2018 and 2019 (Figure S7).

### Singapore Strait community and seasonality

The Singapore Strait samples contained a higher proportion of autotrophic and oligotrophic microbes such as the Synechococcales, Nitrosupumilales, marine group II Euryarchaeota (MGII) and SAR11. The most prevalent cyanobacteria found in the Singapore Strait belonged to the order Synechococcales and were most closely related to *Synechococcus* representative of clade II, which has been identified as the dominant type in warm and oligotrophic waters^46^. The phylum Euryoarchaeota found in high abundance in the Singapore Strait is related to Marine Group II, which is widely distributed in the oceanic euphotic zone^47, 48^ and is the dominant planktonic Archaeon at the surface water of the South China Sea^49^. Marine group III Euryarchaeota (MG-III) was also present in the Singapore Strait, but in a lower proportion. This group is known to be prevalent in deep-sea waters, but few studies have reported their presence in the photic zone^50–52^. Other prevalent members of the microbial communities of the Singapore Strait included the Alphaproteobacteria clade SAR11, which has previously been found to dominate heterotrophic bacterial communities in coastal and open ocean environments^53–55^.

The microbial community shift in the Singapore Strait is synchronous with the seasonal reversal of ocean currents between the Java Sea and the South China Sea^13^ suggesting that the advection of microbial communities from different water basins might be causing this shift. During the SW monsoon, currents flow northward from the Java Sea^13^, bringing less saline water into the Singapore Strait due to the high precipitation over Sumatra, Borneo, and Java. The high freshwater input from these islands into the Java Sea is likely also the source of nutrients transported to Singapore at the start of the SW monsoon period (Figure 1). The net northerly water transport during the SW monsoon transports a community with a higher proportion of Archaea to the Singapore Strait, especially the ammonia-oxidizing archaea *Candidatus* Nitrosopelagicus and *Candidatus* Nitrosopumilus, presumably reflecting higher nutrient concentrations and greater dominance of heterotrophic recycling of organic matter. During the NE monsoon, the current direction reverses and water from the South China Sea flows into the Singapore Strait^13^, carrying more saline and more oligotropic water with less terrestrial input into the Singapore Strait. Consequently, all nutrients decrease in concentration except for a small peak around the beginning of the NE monsoon (Figure 1) that is probably driven by local run-off (since this is typically the period with the highest rainfall in Singapore).

While our study showed an increasing trend in nutrient concentration during the SW monsoon, the chlorophyll did not show any trend. Other parts of South East Asia have also been shown to experience seasonal dynamics in the chlorophyll concentration as a result of monsoon systems^8, 10, 11^. The analysis of the ASVs that differ most significantly between the two monsoons suggested that ecotype replacement is likely occurring during the seasonal cycle. This replacement involves ASVs belonging to SAR11 clades, SAR406 and the NS4 and NS5 groups, which have been shown to display seasonal dynamics in temperate waters^56–60^.

Although few eukaryotic ASVs had their abundance increased or were present during either the SW or NE monsoon period, no clear seasonal pattern was observed among eukaryotes. *Micromonas*, the main photosynthetic picoeukaryotes in our dataset, have been reported as the dominant group in the sub-tropical waters of the South China Sea^61^ and off Taiwan^62^. In Taiwan, high abundances of clade B5 were detected during the summer and autumn, with a peak in July when the local hydrographic characteristics were high temperature and irradiance, and oligotrophy^62^. In our study, *Micromonas* sp. clade B5 seem to have their abundance increased during the NE and inter monsoon months (Figure S7), a period when the water of the Singapore Strait tends to be more oligotrophic.

Members of the *Mesodinium* species complex are known to form periodic or recurrent non-toxic blooms (red tides)^63, 64^. These ciliates can photosynthesize by acquiring and maintaining organelles from cryptophyte prey^65, 66^. Cryptophytes are an important component of phytoplankton communities in coastal ecosystems, especially estuaries environments^67, 68^. High abundances of cryptophytes have been associated with either preceding^69^ or co-occurring peaks^70^ of *Mesodinium* in coastal ecosystems. Although *Mesodinium* and the cryptophyte species complex *Geminigera* were among the abundant groups in Singapore strait, we did not observe a *Geminigera* -*Mesodinium* dynamic in our dataset. Also, to our knowledge, red tides have not been reported in the Singapore Strait.

Blooms of *Coscinosdiscus* and *Chaetoceros*, during the NE and inter-monsoon periods, were reported in the Singapore Strait 70 years ago^15^. Although these taxa were among the dominant phytoplankton group that we found in the strait, neither a seasonal trend nor a bloom were observed during the present study. During the 1968 SW monsoon, a bloom of pennate diatoms (Bacillariophyceae) was also reported^15^, and this group is known to be part of the phytoplankton community in the Singapore Strait^14^. Surprisingly, very few reads in our dataset were assigned to Bacillariophyceae, probably due to the fact that the forward primer used displays one mismatch to all Ochrophyta (data not shown), which is confirmed by the fact that this group was detected through its plastid 16S rRNA sequences (Supplementary Data S2). As this is the first long-term study of the protist communities in the waters of Singapore since the early 50’s^15^, it is impossible to known whether either the absence of blooms today or the appearance of blooms in the past are anomalies of the ecosystem.

## Conclusion

In this 18-month long study we successfully captured the environmental characteristics of two Straits with different trophic status. In the more eutrophic Strait, the Johor Strait, large environmental fluctuations were observed throughout the year, yet no recurring seasonal pattern could detected in the microbial community composition. Conversely, the Singapore Strait showed a seasonal variability which might be a result of the different monsoon regimes. Our study suggests that even in the vicinity of the Equator, where irradiance and temperature show little variation, seasonal trends are reflected in the microbial community composition in relation to monsoon alternation. Based on these findings, we argue that longer sustained observations over a broader spatial scale are needed, in equatorial waters, to identify the patterns of microbial community assembly and the longterm trajectories in the seasonal succession of specific taxonomic groups of Bacteria, Archaea and Eukaryota.

## Acknowledgments

We would like to thank the members of the Singapore Laboratory for Integrated Microbial Ecology and DHI explorer crews for help with sampling. We thank *Indigo V Expeditions* for providing sampling equipment, Christaline George, Sandra Kolundžija, Rosalie Chai and Halimah Razali for help with sampling, Daniela Drautz-Moses for technical assistance with Illumina sequencing, and Chen Shuang for nutrient analysis. This study was supported by the National Research Foundation, Prime’s Minister’s Office, Singapore under its Marine Science Research and Development Programme (Awards No. MSRDP-P13).

## Author contributions statement

CC and FML conceived the study. CC, WW and AK collected and processed the samples. CC, FML, WW, DV and AL analyzed the data. CC and FML drafted the manuscript. CC, DV, AL, PM, FML edited the final version of the paper.

## Additional information

### Data availability

Raw sequencing data have been deposited to GenBank under Bioproject number PRJNA497851. Processed data and scripts are available from https://github.com/slimelab/Singapore-metabarcodes.

### Competing interests

The authors declare no competing interests.

## Notes

https://github.com/slimelab/Singapore-metabarcodes

