## Supplementary Figures for "Temporal dynamics of Bacteria, Archaea and protists in equatorial coastal waters"

### 1 Supplementary material

All supplementary material is available at https://github.com/slimelab/Singapore-metabarcodes

#### 1.1 Supplementary Data

Supplementary Data S1: (monsoonpaper_env_data.csv) List of samples collected with environmental parameters.

Supplementary Data S2 (Singapore ASV_table.xlsx). Sheet ASV: List of ASVs with taxonomic affiliation, assignement bootstrap values for eukaryotes, sequence and number of reads in each sample (only samples from four stations,STJ, EC, SBW and PR are used in this paper). Sheet Blast eukaryotes: Summary of BLAST assignments against PR[230] and GenBank for Eukaryota ASVs (nuclear 18S rRNA).

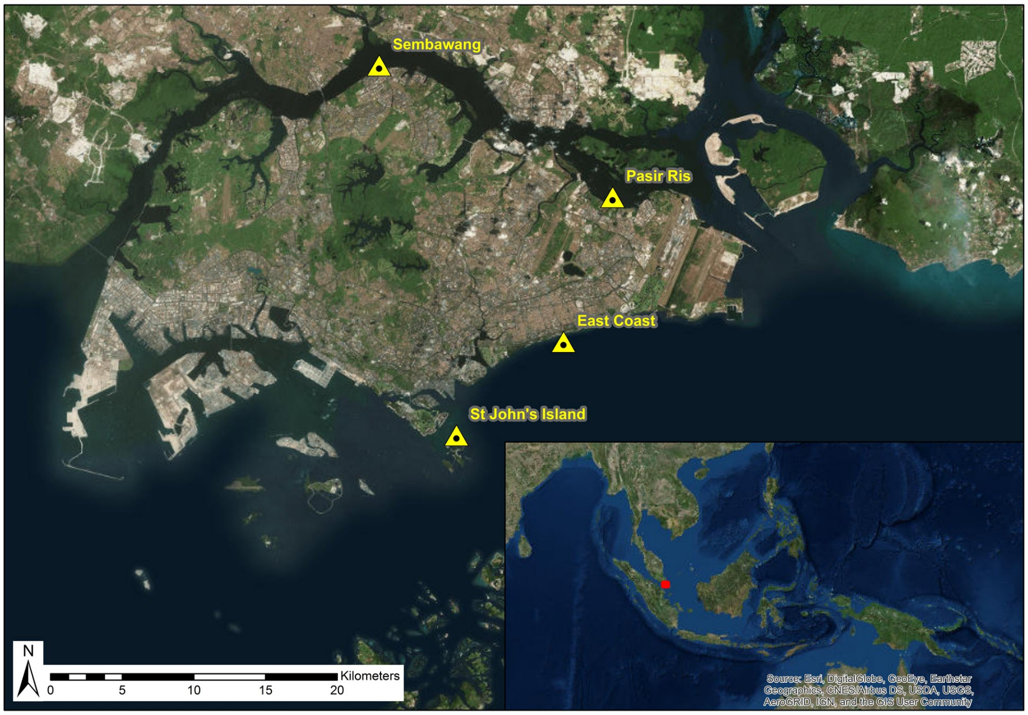


**Figure S1.** Map of Singapore showing the four stations sampled in this study.

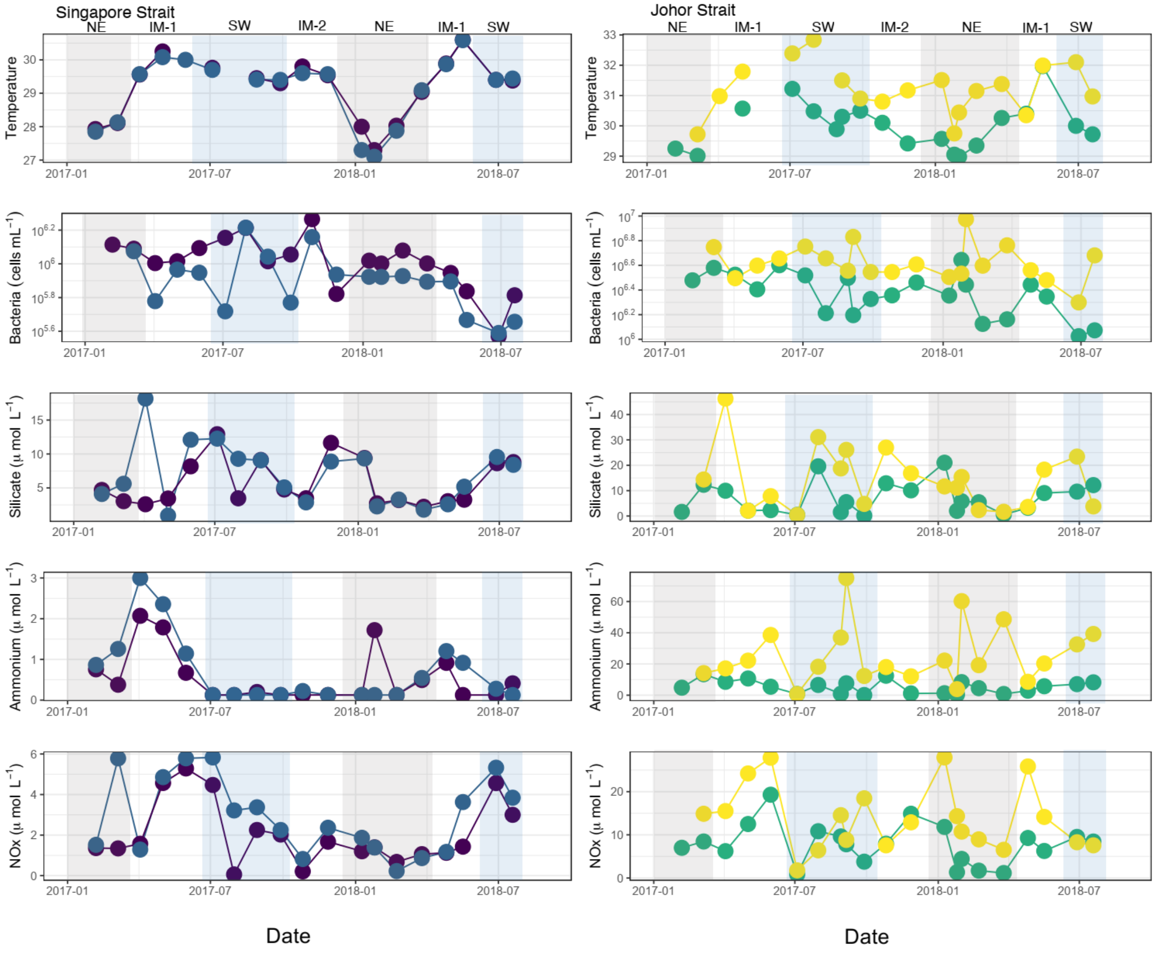


**Figure S2.** Temperature, bacteria abundance, silicates, ammonium and $NO_x$ during the 18-month time series in Singapore coastal waters. Highlights in grey and blue represent NE period and SW periods respectively.

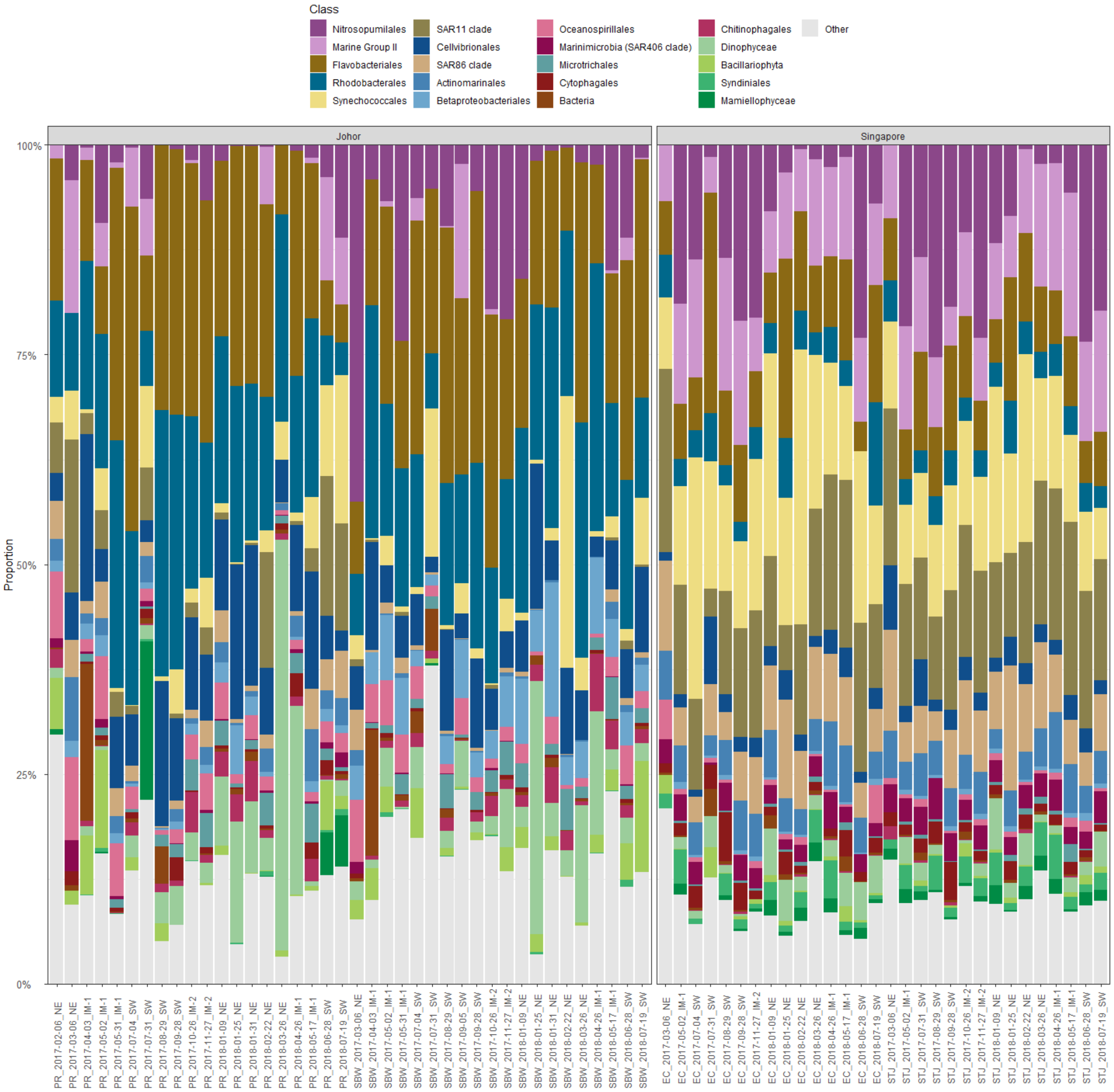


**Figure S3.** Taxonomic composition of the microbial communities in Singapore coastal waters focusing on the top 20 most abundant groups (class level) for the three domains (Archaea, Bacteria, Eukaryota).

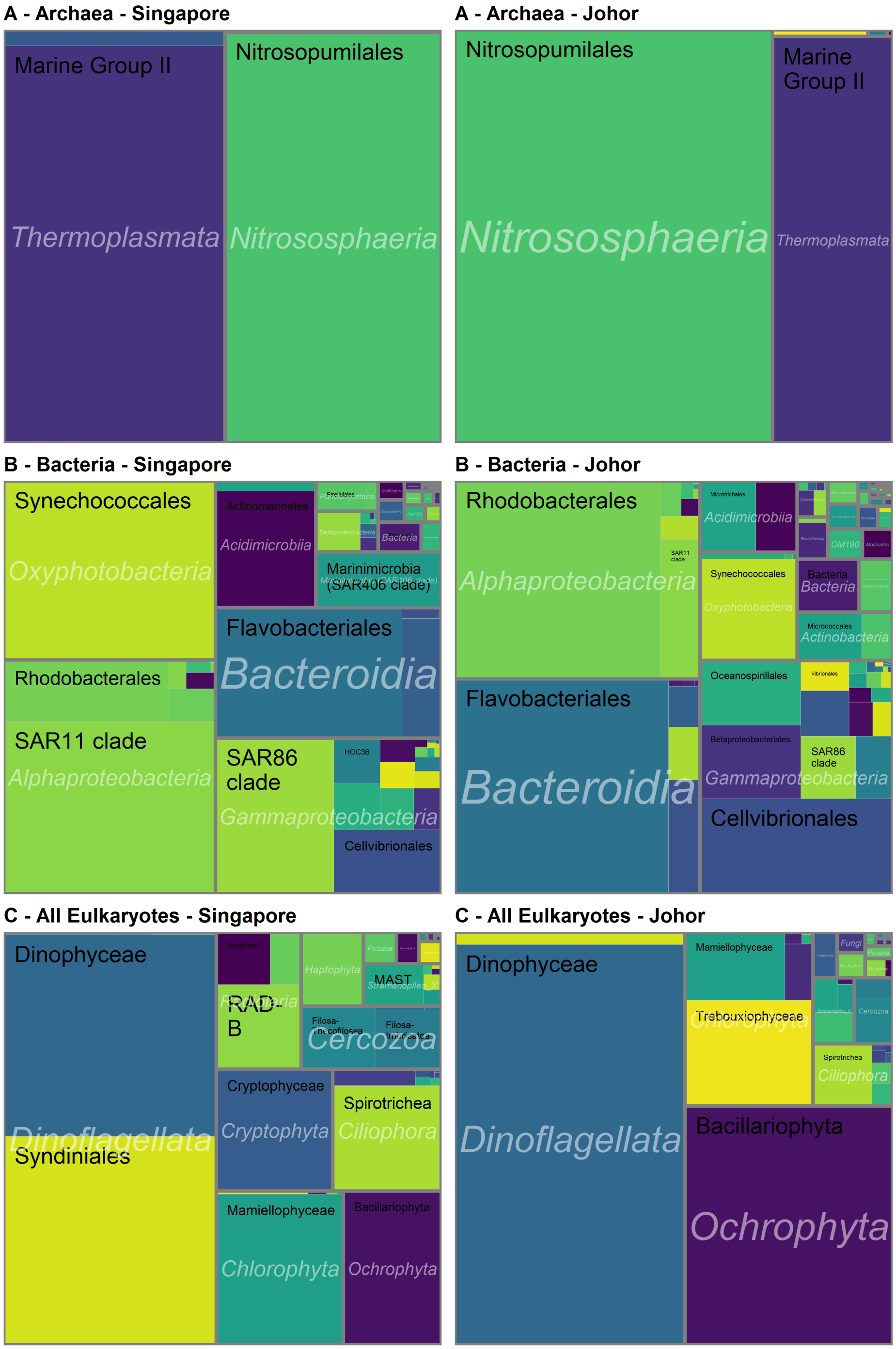


**Figure S4.** Tree map with the major taxonomic groups in Singapore and Johor Straits for the 3 domains: Archaea, Bacteria and Eukaryota.

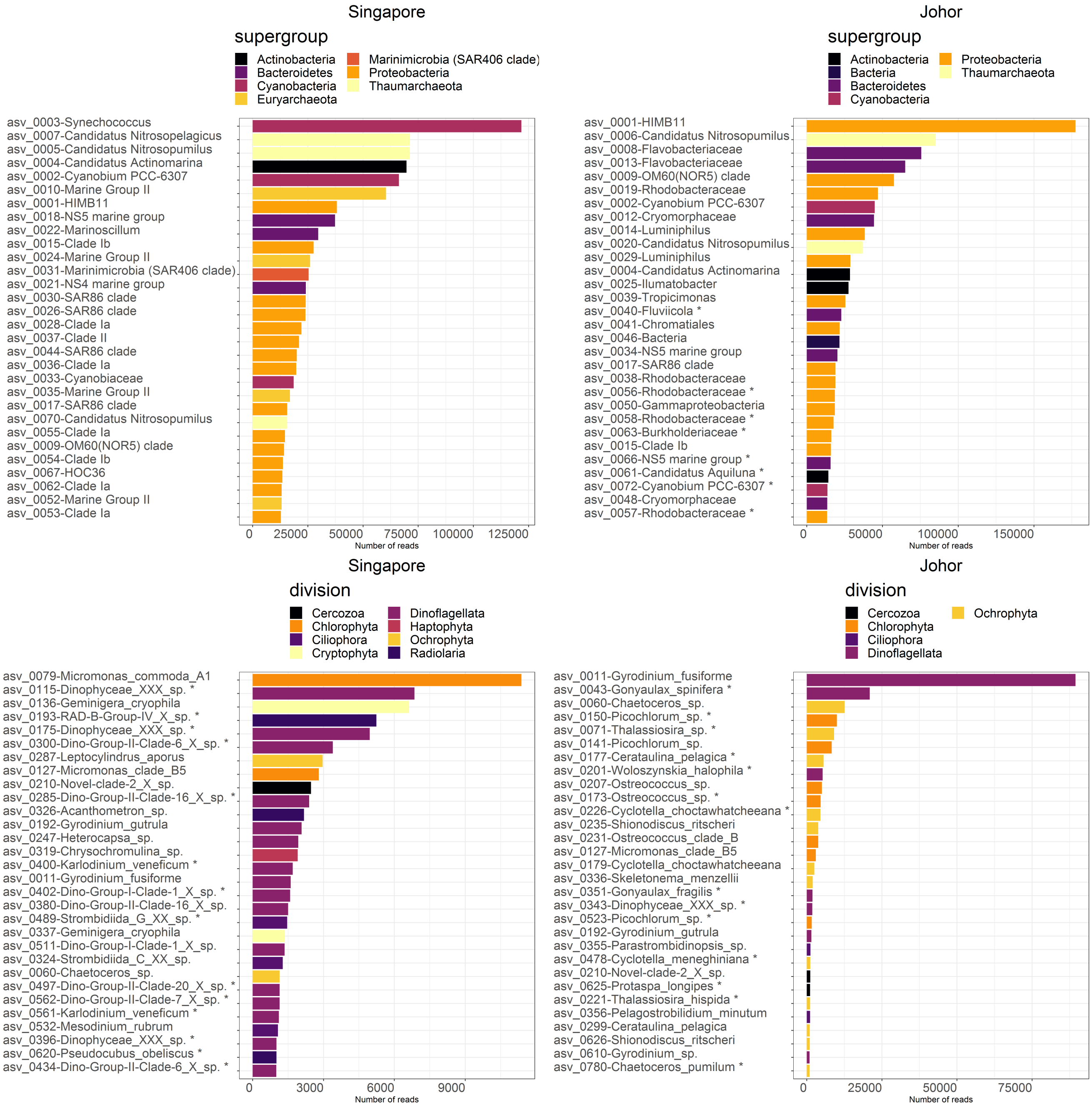


**Figure S5.** Thirty most abundant Archaea and Bacteria (top) and Eukaryota (bottom) ASVs for Singapore (left) and Johor (right) straits. ASVs that are unique to one strait are labelled with an asterisk.

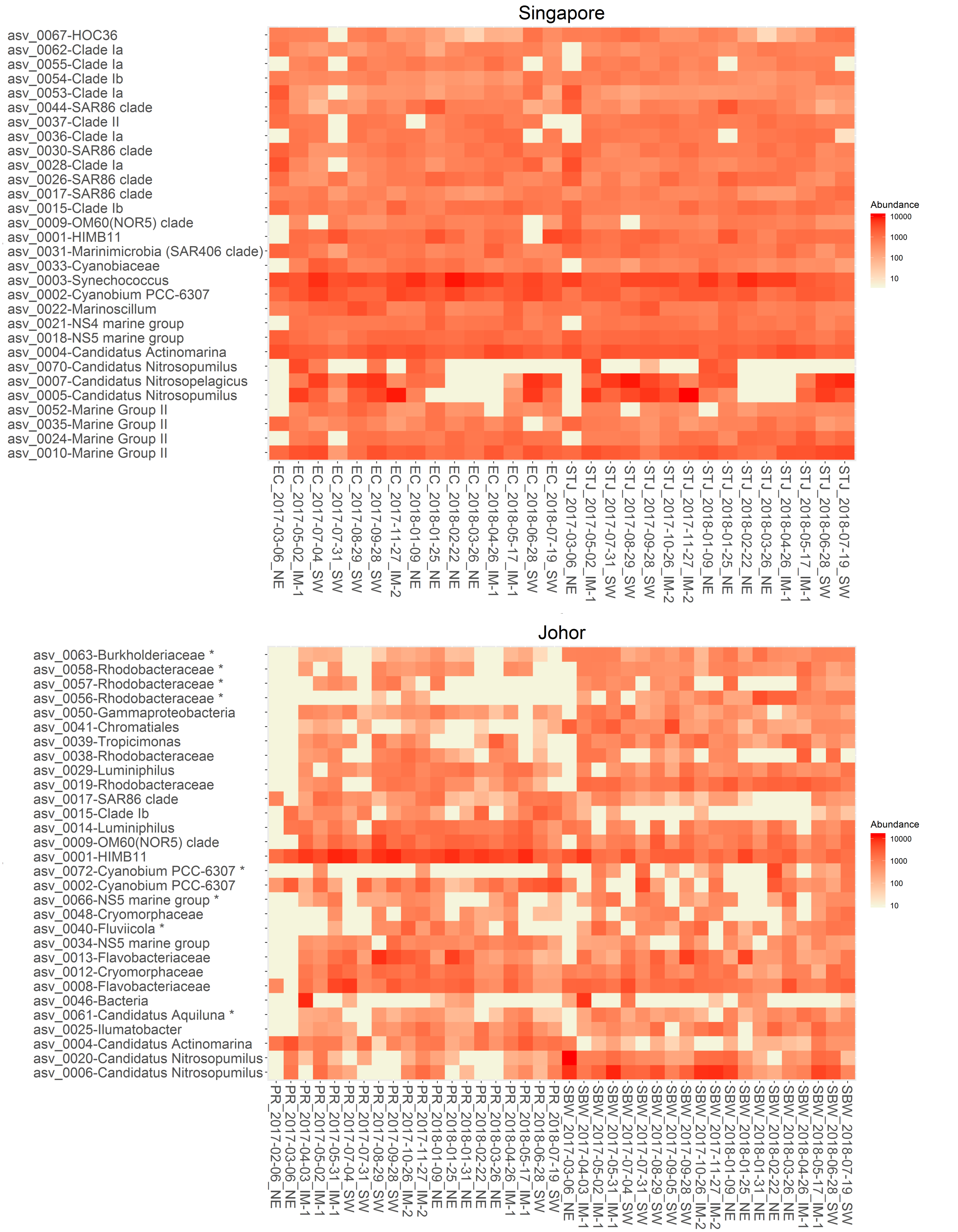


**Figure S6.** Sample heatmap of the thirty most abundant Archaea and Bacteria ASVs for Singapore (top) and Johor (bottom) straits. ASVs that are unique to one strait are labelled with an asterisk.

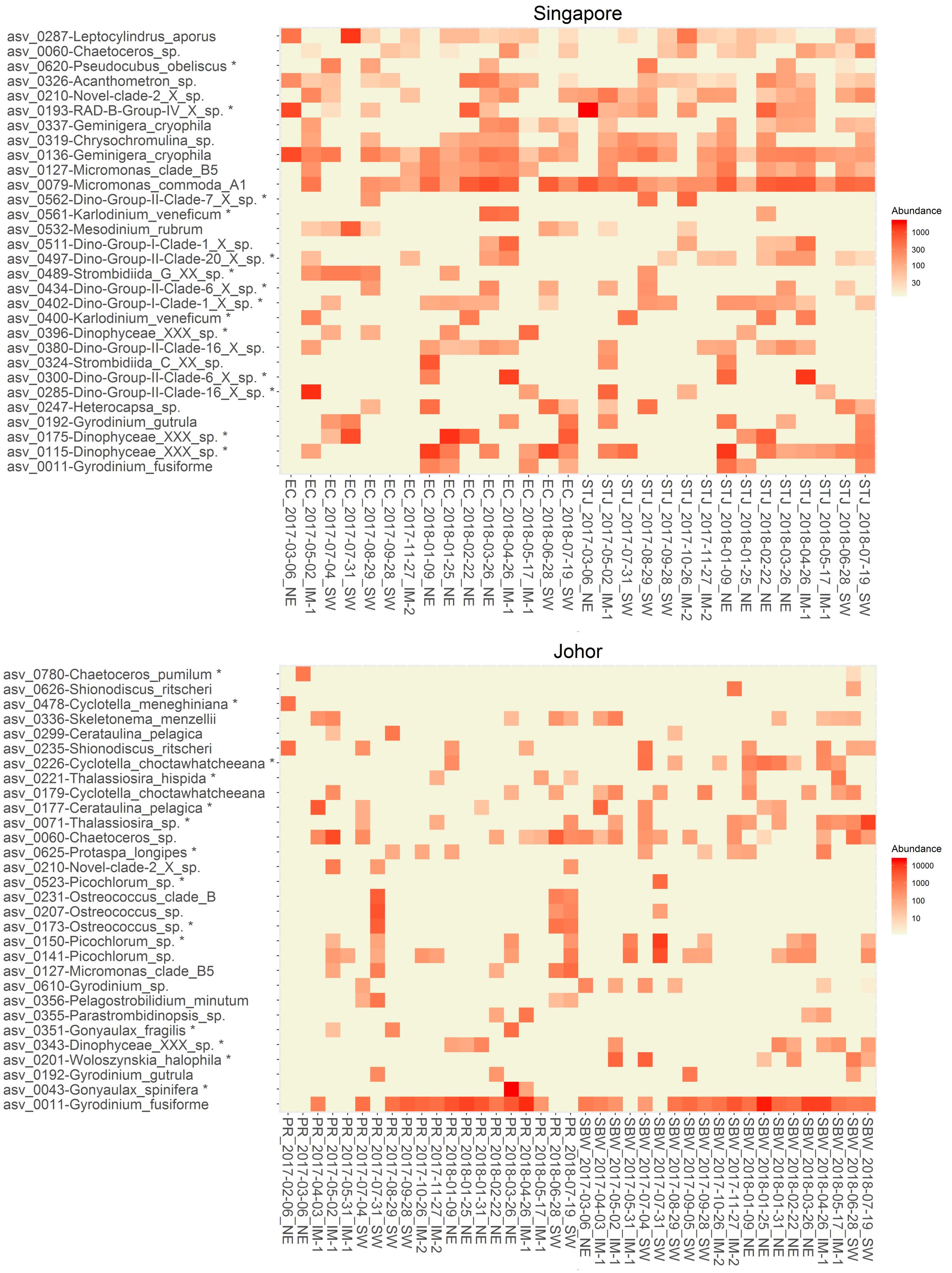


**Figure S7.** Sample heatmap of the thirty most abundant Eukaryota ASVs for Singapore (top) and Johor (bottom) straits. ASVs that are unique to one strait are labelled with an asterisk.

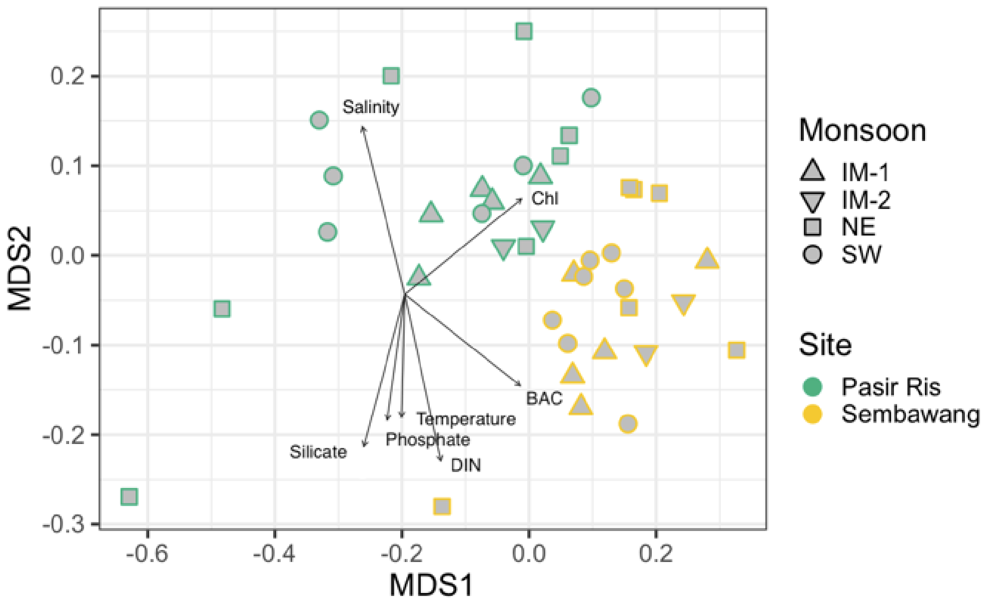


**Figure S8.** Non-metric multidimensional scaling (nMDS) of Bray-Curtis similarity index for Johor Strait (PR and SBW stations). Each sample is labelled based on location and monsoon period. The arrows represent environmental parameters with $p<0.05$ when performing an *envfit* analysis.

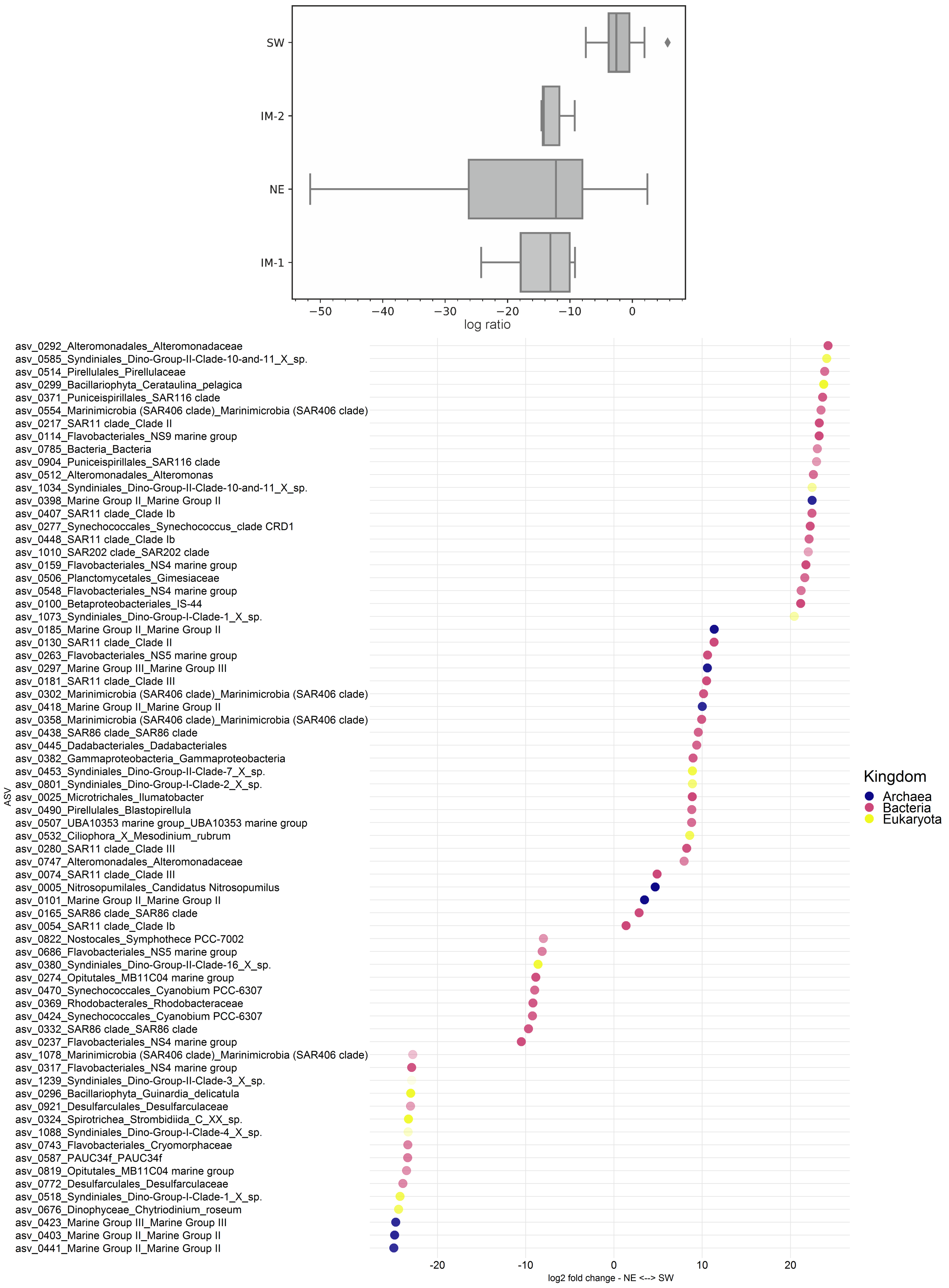


**Figure S9.** Top. Gneiss analysis for Singapore strait based on monsoon period. Bottom. Members of the communities that drive the difference between the NE and SW monsoons based on DESeq2 analysis[33] using a threshold p-value $< 0.01$. Symbol transparency is inversely proportional to ASV rank.

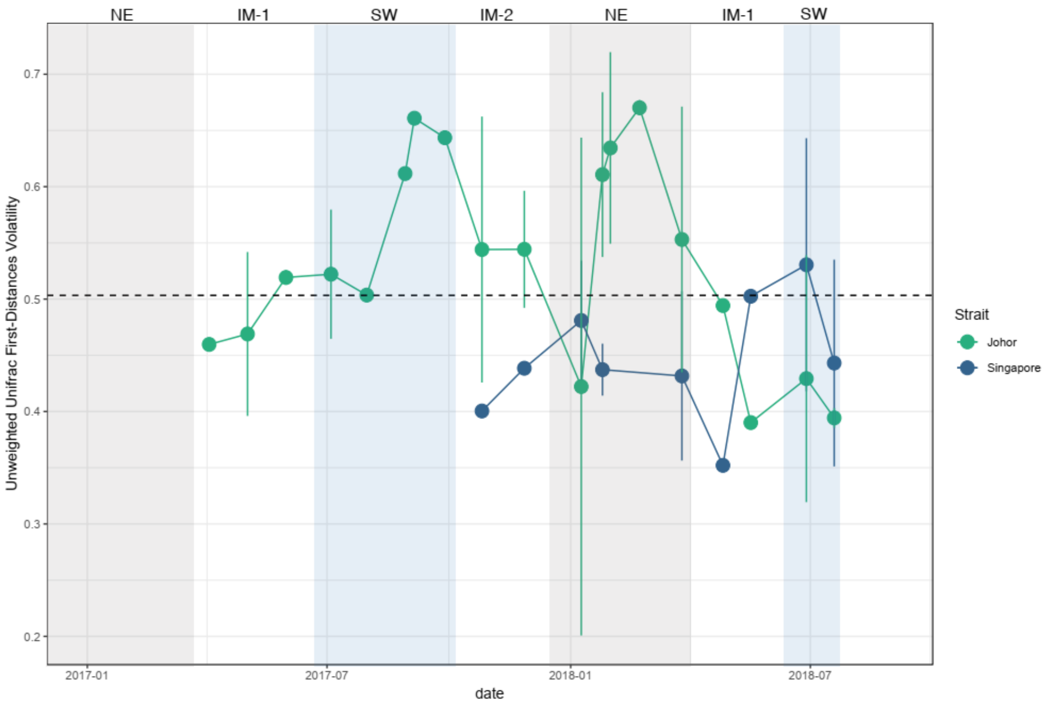


**Figure S10.** First-distances volatility values computed on the unweighted Unifrac distance matrix of the two Straits. The dotted line indicates the global average and error bars the individual sample dispersion from the Strait average

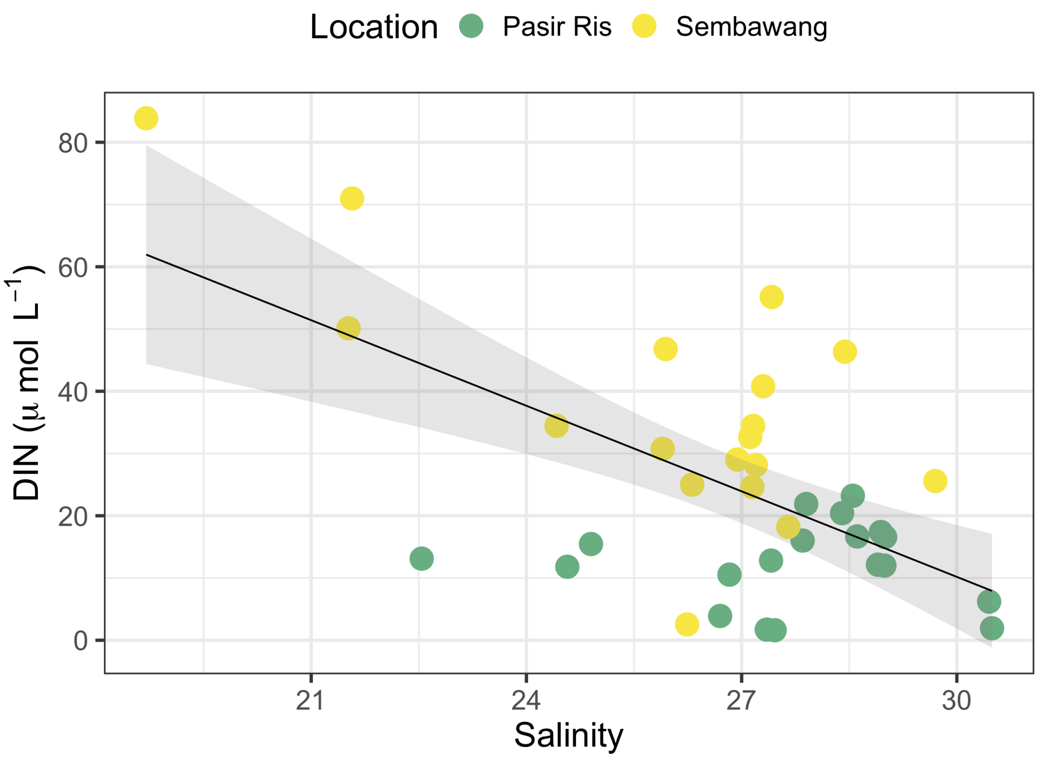


**Figure S11.** Correlation between DIN and salinity in Johor strait.
